# Failure to reproduce the effect of procedural memory interference on wakeful consolidation of episodic memory in younger and older adults

**DOI:** 10.1101/2024.10.17.618844

**Authors:** Lara Kamal, Busra Celik, Griffin Alpert, Samantha Gonzalez, Vian Nguyen, Michael Freedberg

## Abstract

Brown and Robertson (2007) revealed that skill learning interferes with the wakeful consolidation of episodic memories in young adults. This finding is commonly used as evidence that episodic and procedural memories should not be learned in close temporal proximity but has not been reproduced by an independent laboratory. Additionally, older adults experience episodic memory deficits, but it is unknown whether this group is also vulnerable to this type of interference. We aimed to reproduce Brown and Robertson’s (2007) finding in younger adults, while also comparing the magnitude of interference between younger and older adults. Forty younger (18-40 years; n =20) and older adults (≥55 years; n = 20) visited the laboratory in the morning and acquired episodic memories (a list of words) immediately before a procedural finger-tapping (procedural) task. Half of all participants were exposed to a learnable sequential structure. In the afternoon of the same day, participants were asked to recall the episodic memories from the morning session. We found weak evidence of interference for both age groups and no statistical difference in interference between groups. Our results suggest that the interfering effects of these memory types may be negligible or overestimated, and that these memory types can be acquired together without interference.

## Introduction

Episodic memories are created from personally experienced facts, events, or objects [1]. For example, they may include the details of an experienced event, such as the people involved, the location, or the emotions that were experienced. Episodic memory is supported by a distributed network of neocortical association areas and parts of the medial temporal lobe (MTL), such as the parahippocampal cortex and hippocampus [2,3]. Procedural memories, in contrast, are skills and habits acquired through practice [4], riding a bike, and are represented in the motor cortex, dorsolateral prefrontal cortex, cerebellum, and basal nuclei [5–7]. There are two types of procedural memories: motor and cognitive [8]. Motor procedural memory involves movement-related skills, such as riding a bike. Cognitive procedural memory involves mental procedures, like solving a math equation or playing a game with fixed rules, such as the card game Uno™.

Previously, the episodic and procedural memory networks were believed to be isolated systems, as classical lesion studies indicated that damage to the hippocampus, medial temporal lobe (MTL), and parietal cortex specifically impaired episodic memory while leaving procedural memory intact [9–11]. Similarly, dysfunction of the striatum impaired procedural learning while leaving episodic memory intact [11]. Recent functional brain imaging studies, however, show consistent interactions between these networks in neurotypical adults during memory-related tasks [12], and behavioral [13–15] and TMS [16,17] studies confirmed that these interactions are associated with memory performance.

Perhaps one of the most striking findings of behavioral interactions between the two memory types is from Brown and Robertson (2007) who showed that learning a procedural skill immediately after acquiring episodic memories disrupts the consolidation of those episodic memories in younger adults [14]. This work is commonly cited as evidence of competition between the two memory types [18–20], and suggests that the two types of memories should not be acquired in close temporal proximity. This is a non-trivial finding that has implications for how material should be taught in classrooms and how individuals with memory deficits such as older adults [21,22] should approach learning new information. For example, it may be best to allow adequate time between episodic and procedural learning tasks to avoid this type of interference.

The current study has two main goals. First, we aimed to reproduce Brown and Robertson’s (2007) finding, which showed that acquiring a procedural skill immediately after episodic memory formation disrupts the wakeful consolidation of the episodic memories in younger adults. To our knowledge, this finding has been reproduced within the same research group [17,23], but not in an independent laboratory. Thus, we sought to provide the first independent reproduction of this effect. Second, while Brown and Robertson (2007) discovered this effect in a population of young adults (21.1±0.3 years old), it is unknown whether this type of memory interference occurs in older adults and how the magnitude of interference compares between age groups. For our experiment, we formulated two hypotheses: 1) Procedural memory will interfere with the wakeful consolidation of episodic memory in younger (replication of Brown et al., 2007) and older adults; 2) Older adults will show significantly more procedural memory interference on episodic memory than younger adults (AsPredicted.org; #141678). We predicted greater memory interference because older adults undergo substantial neurological changes [24] that can potentially intensify episodic memory deficits and increase vulnerability to procedural memory interference. In younger adults, network organization is highly modular, and the degree of modularity is positively associated with cognitive function [25]. In older adults, however, modularity is reduced [26–28], possibly leading to increased crosstalk between the episodic and procedural memory systems [29] and greater memory interference. Alternatively, these neurological changes might be compensatory, reducing the impact of memory interference. However, there is not enough direct evidence comparing memory interference between younger and older adults. Therefore, replicating Brown and Robertson’s (2007) study with both healthy younger and older adults could offer a more profound insight into the relationship between aging and memory interference, bridging a critical gap in research.

## Methods

### Experimental overview

The study was approved by the University of Texas at Austin Institutional Review Board (#STUDY00000860). Data for this study were collected between 05/03/2023 and 05/16/2024 The objective of this study was to test the reproducibility of Brown and Robertson’s (2007) finding that procedural memory interferes with the wakeful consolidation of episodic memory in younger adults (18-40 years old). Additionally, we included a group of cognitively unimpaired older adults (≥55 years old) to determine whether it also occurs in this group and whether the magnitude of memory interference is different between age groups. In the context of our study, procedural memory interference occurs when it disrupts the wakeful consolidation of episodic memories. Brown and Robertson revealed this interference by showing that acquiring a procedural skill immediately after acquiring episodic memories significantly decreases the retention of those memories compared to a group that performed the procedural task with no learnable structure.

**Figure 1** illustrates the experimental timeline. Participants visited the lab twice in one day (morning and afternoon visits). Before the morning session, participants gave their informed consent. We used the Mini-Mental Status Exam [30] to exclude potential participants with a high likelihood of dementia (defined as a score of <24). Prior to the main tasks, participants completed the Everyday Memory Questionnaire (EMQ; [31]), which is a subjective measure of memory failures in daily life. After completing these measures, participants performed an episodic memory task (memorizing a list of words), followed by the serial reaction time task (SRTT). Six to twelve hours later, participants were invited back to the laboratory and were asked to recall the same words from the morning session without exposure to the list. As in Brown and Robertson (2007), we disambiguated the interfering effects of procedural learning on episodic memory from general forgetting by comparing consolidation between two groups: The “learning” group performed all procedures as described above and the SRTT included a learnable 12-element sequence. In contrast, the “control” group performed the same procedures, but the SRTT did not have a learnable structure (i.e., procedural control task). We pre-registered our study on as-predicted.org (#141678) before conducting our analysis and predicted that while both groups would show significant procedural memory interference on episodic memory consolidation, the magnitude of this interference would be significantly greater in older adults.

**Fig 1.**
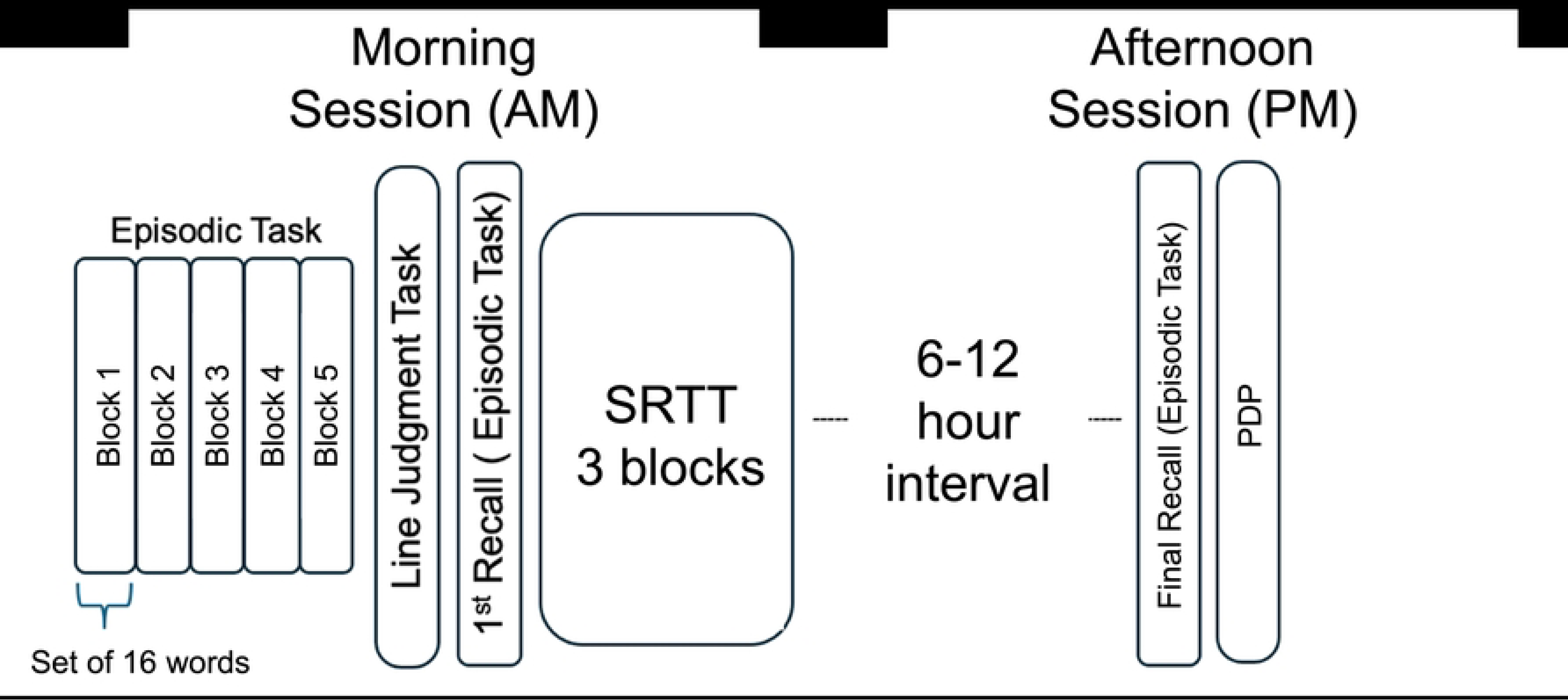
Experimental timeline. SRTT = Serial reaction time task, PDP = Process dissociation procedure

We replicated much of Brown and Robertson’s (2007) experiment, but our study differs in several ways. First, Brown and Robertson (2007) excluded participants who could recall five sequential elements of the procedural task sequence because of previous work showing that offline improvements can be modified by sequence awareness [32]. However, to avoid data removal we included all participants, regardless of sequence awareness and investigated whether explicit sequence awareness affected episodic memory performance, its consolidation across the day, and skill performance after all data were collected. Second, Brown and Robertson (2007) left a period of 10 minutes between the acquisition and recall of episodic memories during the morning session. However, based on the report, it is unclear what participants did during this time frame. To decrease the chances of rehearsal or the chance of any potential differences in the engagement of rehearsal between older and younger adults during this 10-minute period, participants in our study performed a simple behavioral task that minimizes learning demands during this period. Third, whereas Brown and Robertson (2007) assessed sequence awareness through interview, we used the process dissociation procedure (PDP; [33]). The PDP is a structured method for assessing conscious control over sequential information. The PDP is predicated on the idea that if participants have gained conscious control of the sequence, and thus are able to consciously manipulate it, they should be able to recall (conscious control) and avoid (intentional control) the correct element when given the previous elements of the sequence. Lastly, whereas Brown and Robertson (2007) conducted a two-factor ANOVA to test their hypotheses, we used linear mixed effects analyses to separate effects attributable to inter-individual differences from our effects of interest.

### Participants

All participants gave their verbal consent to participate in the study. We recruited 40 participants, consisting of 20 cognitively unimpaired young adults (22.9±3.28 years old) and 20 healthy older adults (65.6±6.97 years old). In total, we tested four groups of ten participants: 1) Younger adults who performed the SRTT learning task (Young learning); 2) Younger adults who performed the SRTT control task (Young control); 3) Older adults who performed the SRTT learning task (Old learning); 4) Older adults who performed the SRTT control task (Old control). Our sample size was kept similar to Brown and Robertson (2007) to facilitate a direct comparison between studies.

Brown and Robertson (2007) showed that procedural memory interferes with the consolidation of episodic memories in healthy younger adults. While the SRTT control group with no learnable structure experienced minimal change in episodic memory performance between visits (M = 0.0, SEM = 0.4), the SRTT learning group experienced a drop in performance between sessions (M = −1.6, SEM = 0.3). This translates to an effect size of 1.43 (Cohen’s *d*). Assuming an alpha of 0.05 and power equal to 0.85, we would have an 85.6% chance of observing a significant memory interference effect in the younger adult age group with a sample size of 10 participants per sub-group (learning and control) using an unpaired, two-tailed t-test. The study was approved by the UT Austin Institutional Review Board and was performed in accordance with their guidelines and regulations. All participants provided their informed consent. All 40 participants were right-handed (Edinburgh Handedness score of ≥ 50; [34]) had corrected-to-normal vision, and were willing to participate. In general, no participants were excluded in our study.

### Procedures

#### Episodic Memory Task

The episodic memory task was performed during the morning session, which took place between 7am and 12pm. During the acquisition phase, participants were instructed to memorize 16 words individually displayed on a computer screen for 2 seconds: Age, Body, College, Door, Figure, Head, Keep, Level, Music, Night, Office, People, Room, Surface, Town, and Week.

Once all the words were presented, participants were prompted to verbally recall as many words as they could, in any order. No time limit was imposed. This block was repeated four more times to facilitate episodic memory acquisition. Following the five training blocks, participants performed a line discrimination task for 10 minutes, during which they compared the length of two vertical lines shown successively on the center of a computer screen. This task was used to prevent mental rehearsal of the word list before recall of the 16 words was tested. Following the line discrimination task, participants were presented with a screen where they were asked to recall the same 16 words without being exposed to the list again. In the afternoon session, participants were once again asked to recall the 16 words for the final time without exposure to the list. No time limit was imposed.

#### Procedural memory task

Immediately following the morning session episodic memory task, participants performed the SRTT. Participants were directed to place the four non-thumb fingers of their right hand on the Chronos stimulus-response device. On each trial, one of four white horizontally aligned open circles (on a black background) would become filled. Participants were instructed to respond as quickly and as accurately as possible to these cues by depressing one of the four spatially compatible responses, where the index (1), middle (2), ring (3), and pinky (4) fingers corresponded with the leftmost to rightmost keys on the button box. Since the Chronos response box includes five keys, one key was taped over, and participants were told to ignore this key. Before starting the main task, participants engaged in a practice block to become familiar with the task where they responded to pseudo-randomly generated cues. The main task during the morning session consisted of three blocks of 280, 400, and 280 trials (960 total). For the learning sub-groups, the first and last 50 trials of each of these blocks included pseudo-randomly generated cues. These were included to reduce the chance of participants developing explicit awareness of the sequence and to compare performance on these trials against performance on sequenced trials. The remaining trials followed a repeating 12-element sequence (2-3-1-4-3-2-4-1-3-4-2-1). Thus, across the three blocks of the morning session, participants in the learning sub-groups performed the sequence 15, 25, and 15 times. The procedural control task was identical to the procedural learning task, except all cues were pseudo-randomly generated.

#### Process dissociation procedure (PDP)

After the final episodic recall, we used the process dissociation procedure (PDP) to assess whether participants gained conscious control over the sequence in the procedural task [33]. This measure was included to explore the potential relationship between the acquisition of explicit sequence knowledge and episodic memory interference and determine how sequence awareness affects the development of a procedural learning in our sample [33,35]. The PDP consisted of two parts. Each part comprised 12 trials, one for each unique triplet in the sequence (e.g., 3-1-2, 3-4-1, 1-3-2, etc.). Participants were cued to respond to the first two elements of the triplet sequence and were then instructed to press (Part 1) or avoid (Part 2) the next correct element of the sequence.

### Data and statistical analysis

Two participants in the Old control group performed the inverse experiment testing the effects of episodic memory interference on procedural skill consolidation six months prior to participating in our study. This experiment included the same word list and the random version of the SRTT as included in our study. Although these participants performed only random keypresses, and thus were not exposed to the same sequence six months prior, they were exposed to the same word list, which could have influenced the results of those participants. However, we included these participants in our analysis based on the six-month gap between experiments and the fact that they did not perform at ceiling on the episodic memory task in the current experiment.

#### Episodic memory interference

Episodic memory performance was assessed during the morning session as recall after the line discrimination task (hereafter referred to as "Ep^1^") and during the second session (Hereafter referred to as "Ep^2^"; **Fig. 1**). We calculated participants change in episodic memory performance as the difference between Ep^2^ and Ep^1^, where higher values indicate stronger consolidation. First, we independently contrasted consolidation scores between learning and control groups for each age group using unpaired, two-tailed t-tests, to test the reproducibility of this finding. Next, we performed a full analysis of our data by submitting episodic memory scores at Ep^1^ and Ep^2^ to a three-factor linear mixed effects model analysis. In the analysis, SUBJECT was included as a random factor and AGE (younger vs. older), SUBGROUP (learning vs. control), TIME (morning vs. afternoon), and their interactions were included as fixed factors. We expected our analysis to yield a significant interaction between SUBGROUP and TIME and a significant three-way interaction between SUBGROUP, TIME, and AGE. Finally, to determine whether initial recall scores at Ep^1^ were related to consolidation scores, we performed a linear mixed effects model analysis of consolidation scores using SUBJECT as a random factor, and Ep^1^ scores, AGE, SUBGROUP, and their interactions as fixed effects.

Our results revealed two null findings: 1) no evidence of memory interference in either group, and 2) no evidence of a difference in the magnitude of memory interference between age groups. These null results raise the possibility that our study was not sufficiently powered to observe these effects. Our rationale for using 10 participants per group was to facilitate a direct comparison between our results and the original report that also included 10 participants per group [14]. Note that although ninety-two participants data were used in Brown and Robertson’s 2007 paper, only twenty participants (10 per group) were included to show that procedural memory acquisition impairs the wakeful consolidation of episodic memories. As mentioned above, our power analysis based on these twenty participants revealed that we would have an 85.6% chance of observing a significant memory interference effect in younger adults with a sample size of 10 participants per sub-group (learning and control) using an unpaired, two-tailed t-test. However, while this power analysis indicates that we are powered to observe an interfering effect of procedural memory acquisition on episodic memory consolidation in younger adults, it does not necessarily indicate that we are powered to observe a significant interaction between memory interference and age group (younger vs. older). Because no studies to our knowledge have examined the effects of procedural memory acquisition on the wakeful consolidation of episodic memories in older adults, we performed post-hoc assessments of power from our effect sizes to determine the number of participants necessary to observe a significant TIME by SUBGROUP interaction (indicating memory interference in both groups) and SUBGROUP, TIME, and AGE interaction (indicating greater memory interference in one age group over the other) using G power [36]. The rationale behind this approach was to determine whether we should continue to collect more participants and retest our hypothesis with a lower alpha or whether this would require an impractical number of participants (e.g., n ≥ 1000 per group).

#### Procedural memory score analysis

Procedural learning scores were calculated during the last block of the SRTT, as in Brown and Robertson (2007), by subtracting the average response time of all correctly performed trials within the last 50 sequence trials (the 50 trials before the final random trials) from the average response time of the final 50 random trials that immediately followed [14,37]. This analysis strategy isolates and removes the contribution of motor control improvements from those attributable to skill learning. These scores were forwarded to our correlational analyses to determine whether the magnitude of procedural learning is negatively correlated with episodic memory consolidation, as found in Brown and Robertson’s (2007) study [14].

#### Examining the relationship between sequence awareness and episodic memory performance and consolidation

As mentioned above, participants were not excluded based on their awareness scores. Rather, we investigated whether sequence awareness was associated with 1) procedural learning, 2) episodic memory scores at Ep^2^, and 3) episodic memory consolidation in our sample. To this end, we employed two-tailed Pearson’s correlation tests examining the relationship between sequence awareness scores and these three variables. Sequence awareness scores were calculated by summing the proportion of correct responses from each part of the PDP and subtracting these scores from chance. Since, during the PDP, participants were not allowed to respond with the same response as the second element of the triplet, chance performance on Parts 1 and 2 was 33.33 and 66.67%, respectively. Thus, awareness scores were calculated as “(Part 1 performance - 0.33) + (Part 2 performance – 0.67),” where values can range from −1 to 1, with positive values indicating some degree of control over the sequence and near-zero or negative numbers representing no control.

#### Additional analyses

We conducted a correlational analysis between age and episodic memory consolidation to determine whether memory interference was related to age. We also performed a correlational analysis between EMQ scores and episodic memory consolidation to examine whether self-reported memory failures in daily life are linked with episodic memory performance during Ep^1^ and episodic memory consolidation. Higher EMQ scores indicate less subjective confidence in one’s memory abilities [31]. Finally, we also conducted a two-factor ANOVA analysis to determine whether the time interval between the morning and afternoon sessions were different across AGE (younger and older) and/or SUBGROUP (learning and control).

## Results

### Assessing memory interference separately in younger and older adults

We conducted a t-test between consolidation scores in the learning and control groups in younger adults. We found a trend suggesting that consolidation was greater in the learning group compared to the control group (*t*(17.02) = −1.77, *p* = 0.09, *d* = 0.79). Thus, our results show a trend in the opposite direction of Brown and Robertson (2007): procedural memory acquisition *enhances* the consolidation of episodic memories in younger adults. A determination of sample size using the Cohen’s d score generated from this contrast (0.79) revealed that a sample size of 24 participants per group is necessary to observe a significant *facilitatory* effect of procedural memory acquisition on episodic memory consolidation (⍺ = 0.05, 1-β = 0.85) using an unpaired, one-tailed t-test. We performed the same analysis for older adults. We found no evidence that consolidation was different between groups (*t*(18) = 0.52, *p* = 0.61, *d* = 0.23). A determination of sample size using the Cohen’s *d* score generated from this contrast (0.23) revealed that a sample size of 267 participants per group is necessary to observe a significant *facilitatory* effect of procedural memory acquisition on episodic memory consolidation (⍺ = 0.05, 1-β = 0.85) using an unpaired, one-tailed t-test. These analyses suggest that procedural memory acquisition *facilitates*, rather than interferes with, the consolidation of episodic memories, but only for younger adults, and that a total sample size of 48 subjects would be required to see this effect in younger adults. Nevertheless, these results do not support our hypothesis that procedural memory acquisition interferes with the wakeful consolidation of episodic memories in younger adults, as seen in Brown and Robertson (2007), or in older adults, and that acquiring more participants would not reveal these effects.

#### Contrasting memory interference between older and younger adults

**Figure 2A** shows consolidation scores for all four sub-groups. We performed a linear mixed effects model analysis on episodic memory scores using AGE (younger vs. older), SUBGROUP (learning vs. control), TIME (morning vs. afternoon), and their interactions as fixed factors, while separating these effects from subject-specific effects. Our analysis revealed a significant effect of TIME (*t*(36) = −3.28, *p* < 0.005), and a trend interaction between AGE and TIME (*t*(36) = 1.77, *p* = 0.09). The time effect indicates that episodic memory scores decreased from the morning to the afternoon, regardless of age or sub-group. The trending interaction suggests that older adults experienced a bigger decrease in episodic memory scores than the younger group.

**Fig 2.**
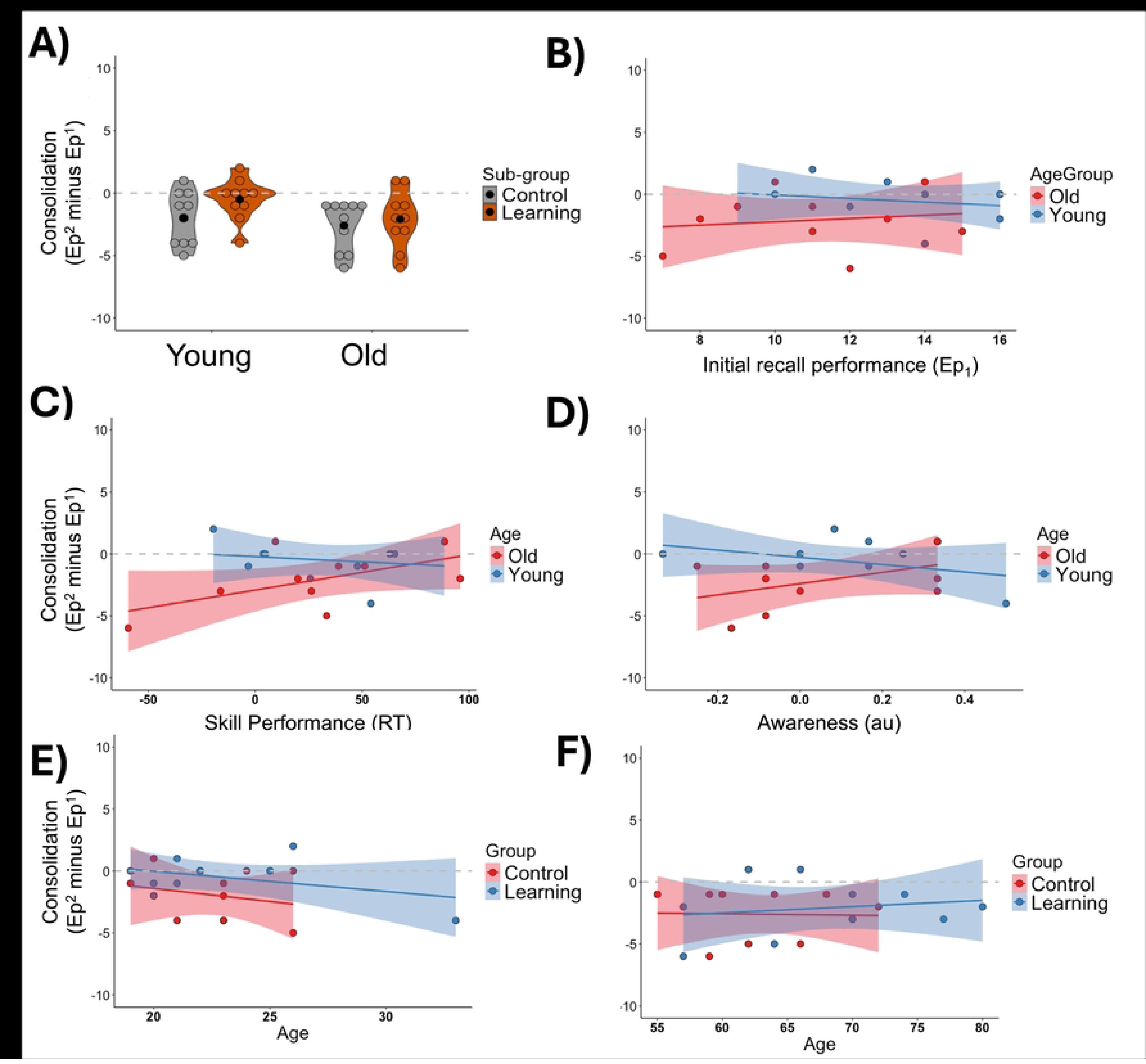
Experimental results. (A) Consolidation scores for both age groups and control (gray) and learning (orange) sub-groups. Black dots represent means. (B-D) Associations between initial recall performance at Ep^1^ and (B) skill performance, (C) sequence awareness, (D) and episodic memory consolidation for younger (blue) and older (red) learning sub-groups. (E-F) Associations between age of younger (E) and older adults (F) and episodic memory consolidation for learning (blue) and control (red) sub-groups.

However, contrary to Brown and Robertson (2007), we found weak evidence of procedural learning interference on episodic memory consolidation across participants of all ages (t(36) = - 0.78, p = 0.44). A determination of sample size using the partial eta squared score generated from an identical ANOVA 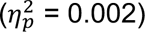 revealed that a sample size of 6,143 participants (∼1,536 per group) is necessary to observe a significant interaction between SUBGROUP, TIME, and AGEGROUP (⍺ = 0.05, 1-β = 0.85). The large number of participants required to see this effect suggests that it is unlikely that our null result was due to a lack of power.

To investigate whether discouraging rehearsal using the line discrimination task decreased episodic memory scores at Ep^1^, potentially affecting consolidation scores, we performed a linear mixed effects model analysis of the effects of AGE and SUBGROUP on consolidation and included Ep^1^ scores as a factor. The analysis revealed no significant effects, including no effect of Ep^1^ scores on consolidation in general (*p*’s ≥ 0.33). Thus, it is unlikely that differences in EP^1^ scores between groups potentially caused by the line discrimination task significantly affected our pattern of results. **Figure 2B** shows the association between Ep^1^ scores and episodic memory consolidation. A direct correlation between Ep^1^ scores and episodic memory consolidation did not yield a significant correlation for younger (*t*(8) = −0.65, *p* = 0.53, *r* = −0.22, 95% CI [-0.75 0.47]) or older (*t*(8) = 0.43, *p* = 0.68, *r* = 0.15, 95% CI [-0.53 0.71]) adults.

### Procedural memory and sequence awareness

Procedural skill learning did not differ between the younger and older learning sub-groups (*t*(16.96) = −0.22, *p* = 0.82, *d* = −0.14) and was not significantly associated with episodic memory consolidation in younger (*t*(8) = −0.53, *p* = 0.61, *r* = −0.19, 95% CI [-0.73 0.50]) or older (*t*(8) = 1.98, *p* = 0.08, r = 0.57, *p* = 0.08, 95% CI [-0.09 0.88]) adults (**Fig. 2C**). Sequence awareness did not differ between younger and older learning sub-groups (*t*(18) = −0.08, *p* = 0.94, *d* = 0.04) and was not significantly related to procedural skill learning [Young: (*t*(8) = 0.51, *p* = 0.62, *r* = 0.18, 95% CI [-0.51 0.73]; Old: [*t*(8) = 0.41, *p* = 0.70, *r* = 0.14, 95% CI [-0.54 0.71]], episodic memory performance at Ep^2^ [Young: [*t*(8) = −0.94, *p* = 0.37, *r* = −0.32, 95% CI [-0.79 0.39]; Old [*t*(8) = 0.80, *p* = 0.45, *r* = 0.27, 95% CI [-0.43 0.77]], or episodic memory consolidation (Young: [t(8) = −1.22, *p* = 0.26, *r* = −0.39, 95% CI [-0.82 0.31]]; Old [*t*(8) = 1.52, *p* = 0.17, *r* = 0.47, 95% CI [-0.22 0.85]]; **Fig. 2D**). These results indicate that it is unlikely that the development of explicit sequence knowledge affected episodic memory consolidation or any potential differences in consolidation between age groups.

### Additional Analyses

We did not find a significant association between age and episodic memory consolidation for the younger (*t*(8) = −1.28, *p* = 0.24, *r* = −0.41, 95% CI [-0.83 0.29]) and older (*t*(8) = 0.51, *p* = 0.63, *r* = 0.18, 95% CI [-0.51 0.73) learning sub-groups (**Fig. 2E and F**). We did not find a significant association between EMQ scores and episodic memory consolidation for the younger (*t*(8) = 0.20, *p* = 0.85, *r* = 0.07, 95% CI [-0.59 0.67]) and older (*t*(8) = −0.17, *p* = 0.87, *r* = −0.06, 95% CI [-0.66 0.59]) learning sub-groups. Finally, we did not observe a significant AGE effect, SUBGROUP effect, or interaction between SUBGROUP and AGE interaction when contrasting the average time interval between the morning and afternoon sessions (*p*’s ≤ 0.22). The average time interval between the morning session and afternoon session was 7.18±0.79 hours (Young = 7.05±0.45 hours; Old = 7.26±0.98 hours).

## Discussion

We tested the reproducibility of Brown and Robertson’s (2007) experiment, which showed that procedural skill acquisition disrupts episodic memory consolidation in younger adults, while also including a group of older adults to investigate whether older adults experience this effect as well and whether there are differences in procedural memory interference across age groups. In our experiment, procedural memory interference was defined as reduced episodic memory performance caused by the acquisition of a procedural skill. We pre-registered our study hypotheses (aspredicted.org #141678), expecting that procedural memory would interfere with the wakeful consolidation of episodic memory in both younger (replicating Brown et al., 2007) and older adults, and that older adults would exhibit significantly greater interference than younger adults. To increase the rigor of our work, we investigated the association between procedural task sequence awareness and episodic memory consolidation to determine whether it is a potential confound. We assessed the conscious control of sequence awareness using the PDP, which is a structured and unbiased measure of the controllability of information. We powered our experiment on the results of Brown and Robertson (2007) and used a linear mixed effects analysis to conduct a more targeted analysis that sequesters the influence of inter-individual variability from our fixed effects of interest. Finally, we discouraged the influence of rehearsal on episodic memory recall during the morning session to prevent potential differences in the benefits of rehearsal on recall between groups. Despite these improvements, we found weak evidence supporting our predictions.

Contrary to Brown and Robertson (2007), our results indicated no significant difference in episodic consolidation between the learning and control groups across morning and afternoon visits for younger or older adults. In fact, our results showed a trend increase in consolidation caused by procedural skill acquisition. Our post-hoc analysis indicated that while adding fourteen more participants to each group may reveal a facilitatory effect of procedural memory acquisition on episodic memory performance, hundreds of older adults would be required to see the same facilitatory effect in older adults. Given this analysis and the fact that Brown and Robertson (2007) reported their effect with the same sample size we used, we are confident that our study was not underpowered. However, our results did not align with theirs. This inconsistency raises the possibility that the effect size we relied on to power our study was too large, or that this effect size is sensitive to the changes we made in our experiment. However, our experimental changes were meant to increase the rigor and sensitivity of our experiment, and it is unclear why these methodological differences would have obscured seeing this effect.

It is possible that discouraging rehearsal may have significantly inhibited the development of episodic memories at initial recall and that this affected our pattern of results. Indeed, comparing recall scores at initial testing between our experiments, participants in Brown and Robertson’s (2007) study acquired roughly 14.5 words at initial testing (learning = ∼15, n=10; control = ∼14; n=10), whereas in our study, participants acquired 12.05 words at initial testing (young learning = 12.00±1.83; young control = 12.60±2.56; old learning = 10.10±2.69; old control = 11.00±2.58). This suggests that differences in the acquisition of episodic memories between studies could explain the discrepancies in our results. However, our control analysis using Ep^1^ scores as a factor did not reveal any evidence that initial episodic memory performance was associated with consolidation. Thus, it is unlikely that the line discrimination can explain the difference in results between our study and Brown and Robertson’s (2007).

Our results do not support the hypothesis that older adults experience greater procedural memory interference on episodic memory compared to younger adults. There are two plausible explanations for our findings: 1) either procedural interference does not affect episodic memory in general, in which case both age groups would exhibit no interference, or 2) there is no discernible difference in interference between older and younger adults. A post-hoc analysis using the effect size from the TIME x AGE x SUBGROUP interaction revealed that it would take thousands of participants to observe this interaction in a follow-up study. Thus, it is unlikely that this null finding was due to a lack of power in our experiment.

Our approach differed from that of Brown and Roberson (2007) in the way we measured participant awareness. In our study, we followed a similar methodology but included all participants regardless of their awareness levels. Brown and Robertson (2007) excluded participants who revealed knowledge of five or more sequential elements of the sequence to prevent the acquisition of explicit knowledge from decreasing skill consolidation. This was most likely used in their investigation of the effects of *episodic memory* interference on *procedural memory consolidation* in their report, but it is unclear whether this exclusion was also performed for the inverse experiment [14], which we replicated in the current experiment. Regardless, we chose to investigate the role of explicit sequence awareness on episodic memory consolidation in our study and found no significant evidence that sequence awareness was associated with episodic memory consolidation, episodic performance on the second visit, or that it was related to skill acquisition in our sample. Nor was awareness significantly different between any group. Thus, we are confident that the development of conscious control over the procedural sequence played a minimal role in shaping our experimental results.

One limitation of our study is that we did not strictly adhere to the 12-hour interval between sessions, as performed by Brown and Robertson (2007). Instead, we used a range of 6-12 hours, which could have impacted our pattern of results. This was performed to accommodate the schedules of participants in our study. On average, our participants returned to the laboratory seven hours after the morning session. Thus, it is possible that if our participants had returned later in the day that we would have observed stronger evidence of procedural memory interference on episodic memory consolidation. Another limitation is that our sample of younger adults (18-40 years old; 22.9±3.28 years) was broader than in Brown and Robertson’s (2007) study (21.1±0.3 years). Although we found no evidence that age was associated memory interference, it is possible that including a broader sample contributed to our null result.

## Conclusion

Brown and Robertson (2007) found that acquiring a procedural skill immediately after an episodic task disrupts the consolidation of episodic memories in young adults, and vice versa. However, this phenomenon has not been reproduced by an independent research group or studied in older adults. To address these gaps, our study replicated Brown and Robertson’s (2007) experiment while including older adults, aiming to both reproduce their findings and investigate the effects of this interference in an older population. We followed a similar methodology but improved the analysis in several aspects. Our results revealed no significant evidence of procedural interference on episodic memory in younger or older groups and we did not observe a difference in the magnitude of interference between age groups, despite using a more appropriate statistical analysis (linear mixed-effects modeling) and ensuring our study was adequately powered based on the original report. Our results suggest that the interfering effects of these memory types may be negligible or overestimated, and that episodic and procedural memories can be acquired together without interference.

